# Mapping the 3D genome through high speed single-molecule tracking of functional transcription factors in single living cells

**DOI:** 10.1101/568675

**Authors:** Adam J. M. Wollman, Erik G. Hedlund, Sviatlana Shashkova, Mark C. Leake

**Affiliations:** Biological Physical Science Institute, Departments of Physics and Biology, University of York, YO10 5DD York, UK

**Keywords:** 3D, yeast genome, transcription factors, single-molecule, transcription, gene regulation

## Abstract

How genomic DNA is organized within the cell nucleus is a long-standing question. We describe a single-molecule bioimaging method with super-resolution localization precision and very rapid millisecond temporal resolution, coupled to fully quantitative image analysis tools, to help to determine genome organization and dynamics using budding yeast *Saccharomyces cerevisiae* as a model eukaryotic organism. We utilize astigmatism imaging, a robust technique that enables extraction of 3D position data, on genomically encoded fluorescent protein reporters that bind to DNA. Our relatively straightforward method enables snapshot reconstructions of 3D architectures of single genome conformations directly in single functional living cells.

## 1. Introduction

Variations in 3D genome architecture contribute to a large number of disorders, including autism, schizophrenia, congenital heart disease and cancer [1]. However, our knowledge of the functional, organization of dynamic DNA in the complex, crowded physiological milieu of living cells remains limited. New methods to elucidate 3D genome structure may be valuable in improving our understanding of not only the native genomic architecture in normal cells but also of the development and progression of diseases associated with DNA structural abnormalities.

There are various existing tools to study 3D genome configuration, such as probing RNA-chromatin interactions, chromosome conformation capture (3C) techniques and microscopy-based approaches, including the 3C variant Hi-C that extends the capability of the technology by identifying longer range interactions across the whole genome [2], [3]. However, none of these methods are comprehensive on their own in regards to generating data representing a dynamic structure of an individual genome conformation from single, functional, living cells [4]. For example, 3C variant techniques are genome-wide, but the results represent the ensemble average of all genome configurations, and so lose dynamic information. Moreover, these methods cannot be performed *in vivo* and, furthermore, are population level techniques generating information from often several thousands of cells and so struggle to render important information concerning cell-to-cell variability, arguably a key feature in ensuring cell survival during conditions of high stress. Standard fluorescence *in situ* hybridization, FISH, is a traditional microscopy-based approach, which is widely used in DNA localization studies. 3D-FISH in combination with confocal microscopy and image reconstruction enables the analysis of the spatial arrangement of chromosomes. However, this technique, in its traditional form at least, requires sample fixation [5], and thus fails to render information concerning structural fluctuations in the genome with time. Recent advances in single-molecule fluorescence microscopy have provided fundamental insights into the interactions of proteins with DNA upon gene regulation in both prokaryotes and eukaryotes [3], [4].

Studies on live cells from a range of different species show that several types of proteins which bind to DNA, including those involved in chromatin remodeling, DNA replication, transcription and repair, operate as oligomeric clusters [6]–[9].

Here we describe a novel approach for achieving dynamic 3D spatial resolution at millisecond time scales and single-molecule detection sensitivity directly in single living eukaryotic cells using astigmatism imaging [10]. We modified a method that generates a narrow field of laser illumination which produces high excitation intensities in the vicinity of single live cells [11]–[14]. This technique is based on introducing astigmatism into the imaging path through insertion of a long focal length cylindrical lens between the microscope emission port and camera detector, which enables extraction of 3D spatial positions of single fluorescent reporter molecules. Astigmatism-based approaches allow imaging over an axial range comparable with the length scale of the nucleus in yeast cells. The method is also relatively easy and cheap to implement compared to competing techniques, such as multi focal plane imaging [15] and approaches which use helical shaped point spread function (PSF) excitation profiles [16]. Astigmatism imaging combined with Stochastic Optical Reconstruction Microscopy (STORM) has been used to image microtubules and clathrin coated pits in cells with spatial resolution which is an order of magnitude better than standard diffraction-limited optical resolution. However, STORM requires typically long imaging times so rapid dynamics are largely lost [17]. In a recent review of 3D imaging techniques, astigmatism imaging approaches perform well in lateral and axial resolution, as well as the axial range over which probes can be detected [18]. Multi focal plane imaging, most simply including biplane imaging, and double helix PSF microscopy, perform marginally better in regards to spatial resolution but these modalities are often complex and/or costly to implement, e.g. requiring multiple objective lenses and/or phase modulation optics. Recently, tilted light sheet microscopy combined with PSF engineering was able to map out the whole mammalian cell nuclear envelope [19] and may become a powerful future technique for 3D genome architecture.

We utilize the budding yeast *Saccharomyces cerevisiae* and its DNA-binding Mig1 protein as a reference for genome mapping. Mig1 is a Zn-finger transcription factor which binds to target DNA sequences under glucose-rich extracellular conditions to repress expression of genes essential for metabolism of non-glucose carbon sources [20], [21]. We showed previously that Mig1 molecules with apparent 2D diffusion coefficients lower than ~0.1 μm^2^/s were immobile and likely to be bound to DNA, and that we could use 3C models combined with bioinformatics analysis to predict the likely Mig1 binding sites in 3D [7]. In our present work here, we directly image fluorescent Mig1 in 3D, and identify immobile Mig1 foci. We then compare our observations to the 3C model and provide valuable biological insights into the 3D eukaryotic genome architecture in single living cells.

## 2. Materials and methods

### 2.1. 3D super-resolution single-molecule microscope

We constructed a bespoke astigmatism super-resolution fluorescence microscope, built around the body of a Nikon *Ti*-series epifluorescence microscope. A schematic of the optical design is shown in Fig. 1. We implemented Slimfield illumination to observe single GFP molecules in living cells, a method that generates high laser excitation intensities in the vicinity of single cells thereby enabling millisecond [7]. Vortran 50 mW 473 nm and 561 nm wavelength lasers, coupled together using a dichroic mirror, were incident on a lens in a telescope with the objective lens to generate a collimated ~20 μm (full width at half maximum) beam at the sample, with an intensity of typically 2.5-3 kW/cm^2^. The image was collected by a 300 mm focal length tube lens onto a Photometrics Evolve 512 Delta EMCCD camera, with a DV2 color splitter to enable separate, simultaneous imaging of GFP and mCherry fluorescent protein components in the sample. A cylindrical lens was placed between the tube lens and the camera for astigmatism imaging. The resulting magnification at the sample is 93 nm per pixel. The sample was held on a Mad City Labs XYZ positioning nanostage.

**Figure 1:**
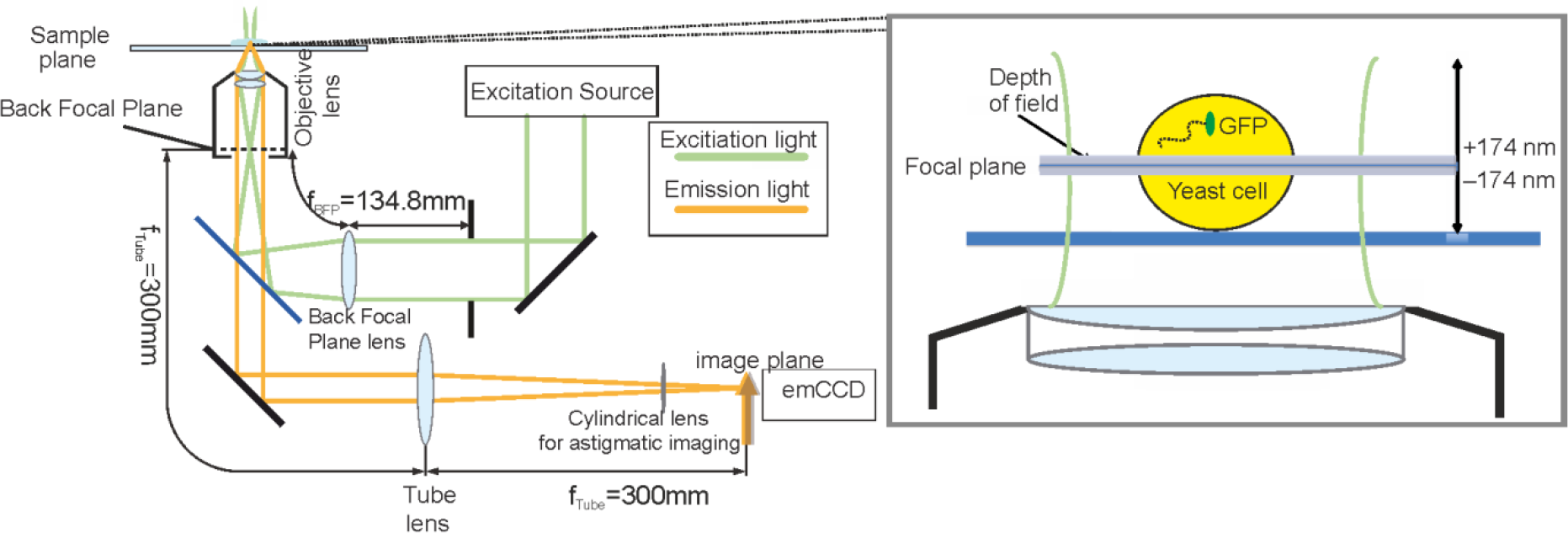
The astigmatism microscopy imaging system. The depth of field in a non-astigmatism microscope is indicated with the shaded band (zoom-in image on the right panel). The cylindrical lens is located just outside the imaging port of the microscope.

### 2.2. Calibration of the microscope

**Figure 2:**
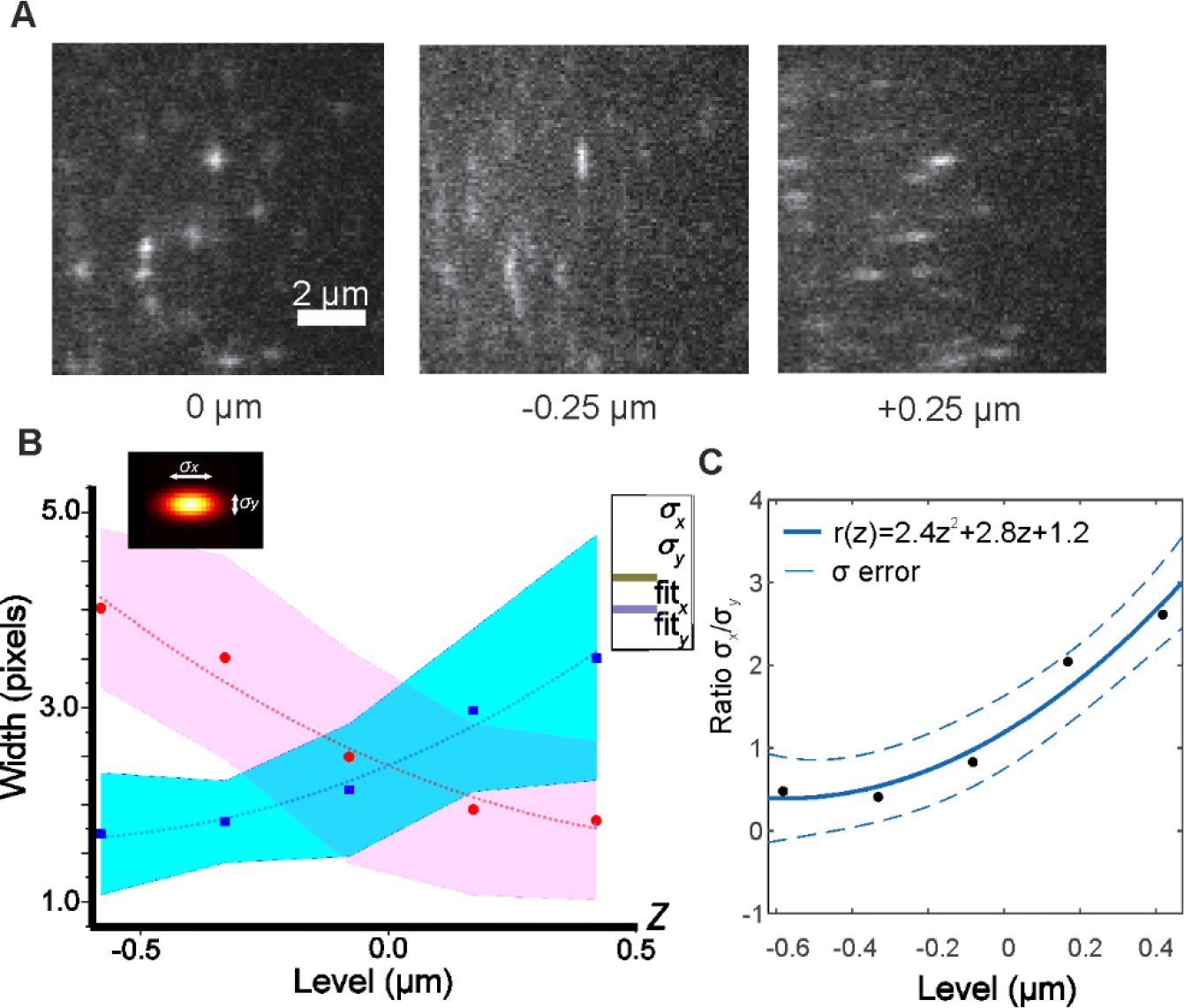
A: Micrographs of in vitro surface-immobilization assay, with purified GFP molecules bound to a glass coverslip surface. Here we illustrate offsets of −0.25 μm, 0 μm, and +0.25 μm to show the deformation of the PSF image at different depths. B. Mean fitted x and y sigma values to in vitro GFP as a function of focal depth (blue and red squares) with standard error indicated as shaded area and 2^nd^ degree polynomial fits as dashed lines. C. The ratio of the fitted Gaussian sigma x to sigma y values as a function of focal depth (black squares) with 2^nd^ degree polynomial fit was used to calculate z position and 1 sigma confidence interval values (full and dashed blue lines, respectively).

#### 2.2.1. Fluorescent protein *in vitro* assay

To calibrate the fluorescent foci PSF image deformation due to the cylindrical lens, we imaged immobilized GFP on a coverslip at different axial distances, using a surface-immobilization assay adapted from earlier studies [22], [23]. In brief, a ~2-3 mm width channel chamber with a volume of ~5 μl was created from two strips of a double-sided tape on a microscopy slide and covered with a plasma-treated BK7 coverslip. A PBS solution of 2 μg/ml anti-GFP antibody (Invitrogen, G10362) was flowed into the chamber and left to adhere to the coverslip surface for 5 min at RT. Excess antibody was washed away by 200 μl of PBS. Four chamber volumes of 1 μg/ml GFP were then injected into the chamber, left to conjugate with antibodies for 10 min, and washed to remove any unbound molecules. 300 nm diameter polystyrene beads (Invitrogen, C37281) in 1:1000 dilution were added to the slide to focus on the coverslip surface in the brightfield, before the nanostage was moved −150 nm to set the *z* = 0 position. Images of single immobile GFP molecules were acquired at a 5 ms exposure time at axial positions between *z* = −0.5 μm and *z* = +0.5 μm in 0.25 μm intervals (Fig. 2A).

#### 2.2.2. Axial distance calibration curve

Images of immobilized GFP molecules were tracked using bespoke software written in MATLAB (MATHWORKS) [24], modified to fit each fluorescent foci PSF image using a standard 2D lateral Gaussian function with independent σ width parameters, σ_x_ and σ_y_. The mean σ_x_ and σ_y_ values were collated for each axial position and are shown in Fig. 2B, indicating similar results to other 3D microscopes [25]. Ultimate calibration of the ratio of sigma widths, *r*=σ_*x*_/σ_*y*_, to the axial position, z, was obtained by fitting an optimized 2^nd^ order polynomial (Fig. 2C):

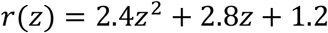

which gave a goodness-of-fit parameter *R*^2^ = 0.96. The data can also be fitted by an explicit de-focusing equation if required [26] but the form of the fit is in practice not critical since model-dependent differences to fits in general are small compared to the actual empirical axial precision [27].

### 2.3. Simulation of 3D tracks

**Figure 3:**
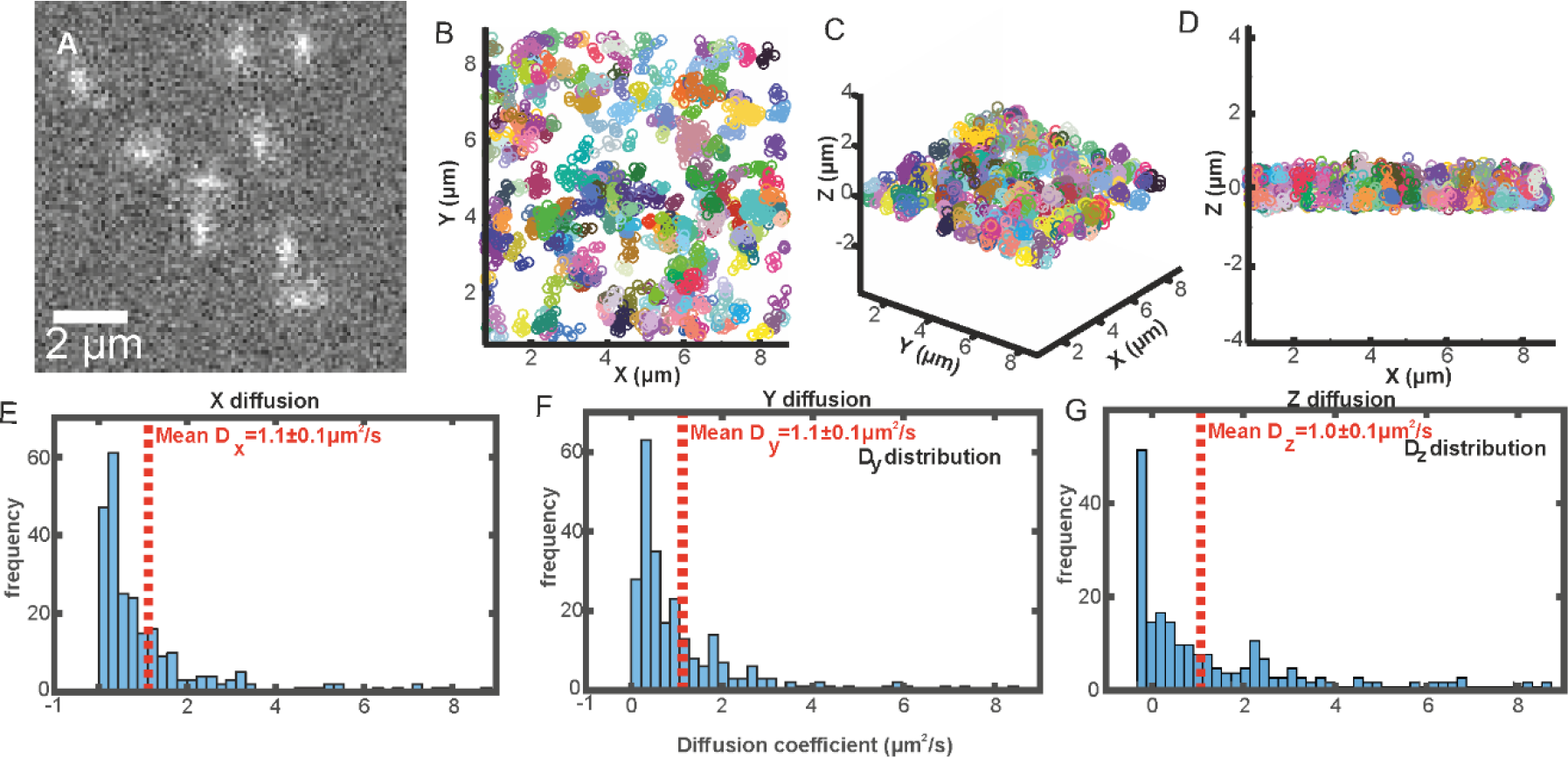
A single 5 ms expore time sample image frame taken from A. simulated sequence of fluorescently labelled molecules diffusing in three spatial dimensions B-D. Scatter plots of tracked simulated diffusing molecules. E-G. Distribution of diffusion coefficients of tracked simulated data in all three spatial dimensions.

In order to verify that our astigmatism microscope could be used to track diffusing molecules as a function of time and verify our calibration, we simulated extended kinetic series of diffusing molecules whose fluorescence intensity was consistent with those measured experimentally, with realistic levels of added noise. We then measured their apparent microscopic diffusion coefficients using exactly the same detection and tracking analysis algorithms as for the experimental data. Astigmatism-deformed fluorescent foci were simulated by taking the average real mean foci image at each axial displacement, then linearly interpolating between each image at intervals of the equivalent camera pixel magnification which was 93 nm per pixel (SI Fig. 1A and B) to create a series of reference images spanning a 1 μm range in *z*, comparable to the approximate axial working range over which we could reliably detect single GFP molecules. 4D Foci positions (i.e. spatial coordinates for *x, y, z,* and also the time dimension *t*) were simulated using Brownian motion with a nominal diffusion coefficient of 1 μm^2^/s based on sensible experimental estimates from earlier 2D measurements [7]. The correct reference image was added to an array at each 4D position, and then realistic camera and signal noise were added to the array (we used a mean camera offset value of 100 counts with Poisson-distributed noise) (Fig. 3A). Images were tracked similarly for the *in vitro* calibration data (Fig. 3B-D), their mean square displacements (MSD) were calculated separately in each dimension (SI Fig. 1C), then their apparent microscopic diffusion coefficients were calculated from a linear fit to the first four MSD time interval values (Fig. 3E-G) using a previously optimized method [28].

### 2.4. Live cell microscopy

For live cell imaging, we used the model unicellular eukaryote of budding yeast *S. cerevisiae*, strain YML14, expressing genomically integrated Mig1-GFP (Mig1 is a transcription factor acting as a repressor for several target genes implicated in glucose metabolism) and Nrd1-mCherry (Nrd1 is a protein component of the RNA polymerase as is a clear marker for the position of the nucleus) fusions [7]. Cells were grown in minimal YNB media (1.7 g/l Yeast Nitrogen Base without amino acid and (NH_4_)_2_SO_4_, 5 g/l (NH_4_)_2_SO_4_, 0.79 g/l complete amino acid supplement as indicated by the manufacturer) supplemented with 4% glucose until mid-logarithmic growth phase, washed and placed into 3 ml of fresh medium for about 1 h. 5 μl of the culture was applied onto a 1% agarose pad perfused with YNB, formed using a 125 μl volume Gene Frame^®^ (Thermo Scientific) and covered with a plasma-cleaned BK7 22 × 50 mm glass coverslip. Typically 1-4 cells per field of view were imaged using conditions similar to those described previously [7], [29].

### 2.5. Analysis of live cell data

Images of Mig1-GFP were tracked using the same methods as before [7]. Frame averages were taken over five consecutive images of Mig1-GFP and Nrd1-mCherry, segmented for the cell and nucleus respectively using the GFP and mCherry signals respectively. These images were then thresholded, which created distinct masks for the nucleus, though often left multiple cells joined together in a single mask. To overcome this issue an extra watershedding step [30] using the nucleus masks as seed basins for the joined cell masks, allowed true cell masks to be obtained of each separate cell. Mig1-GFP foci tracks were then assigned into cells and the separate sub-cellular compartments (i.e. cytoplasm or nucleus) based on their positions. The apparent microscopic diffusion coefficients of each foci track were calculated as in our previous 2D study but now using the full MSD determined from the complete 3D spatial localization data. We used the diffusion coefficient threshold defined in our previous study to collate the putative immobile Mig1 tracks as being those with a rate of diffusion at or below 0.1 μm^2^/s [7]. We then calculated the fluorescence intensity centroid of these tracks to define the position of immobile Mig1 in the nucleus, and thus a putative Mig1 binding site on the genomic DNA.

## 3. Results and discussion

### 3.1. Calibration and performance of the microscope

We estimate that our calibration yields an axial resolution of ~100 nm, roughly 2-3 times poorer than our measured lateral resolution of ~40 nm under comparable imaging conditions [31]. This reduction in spatial resolution from lateral to axial is similar to other previously implemented 3D light microscopes [25] and compares favorably with other astigmatism based microscopes. Although others have reported superior axial resolution using astigmatism approaches, for example Huang [27] achieved 30 nm lateral resolution and 50 nm axial resolution using a 3D astigmatism STORM instrument imagingbright organic dyes. Fluorescent proteins probes are more challenging due to poorer photophysical properties, though Moerner reported axial precisions of ~40 nm using yellow fluorescent protein by employing a double helix PSF method with 30 ms per frame sampling [32]. However, since GFP emits approximately an order of magnitude fewer photons on average than YFP prior to photobleaching, and our method involves much faster sampling, also by close to an order of magnitude compared to Moerner’s YFP study, our reported axial precision is close to expectation based on the effective signal-to-noise ratio [33]. Our calibration using single GFP molecules also in many ways represents a worst case for axial resolution as many of the transcription factors we image are clustered [7]. New fluorescent proteins, such as mNeonGreen, are also ~3x brighter [34] and will also increase the axial resolution by a factor roughly equivalent to the square root of the increase in brightness. Astigmatism also has advantages over double helix PSF microscopy: it is easier to implement in the microscope, it does not require permanent alignment of often expensive phase masks – allowing both standard widefield imaging and astigmatism imaging with a flip-in component, and the data analysis is simple.

Tracking simulated foci trajectories, we were able to measure the same apparent microscopic diffusion coefficients as simulated within expected sampling error in all three spatial dimensions. Thus, we are able to track molecules *in vivo*. Our measured diffusion coefficient distributions are broad but this is expected from the long tail of Gamma shaped probability functions [35]. Also, fitting to MSD *vs*. the time interval parameter to generate diffusion coefficients inherently generates positively skewed distributions as extreme high values are not detected due to the limits of tracking while errant low diffusion coefficients result from poor tracking. The latter is due to linking different foci incorrectly into the same trajectory, resulting in apparent reduction in MSD and fits which tend towards low values. Both these factors broaden the measured diffusion coefficient distributions, but our simulations show that the correct population statistics can still be extracted.

### 3.2. 3D architecture of Mig1 binding sites

**Figure 4:**
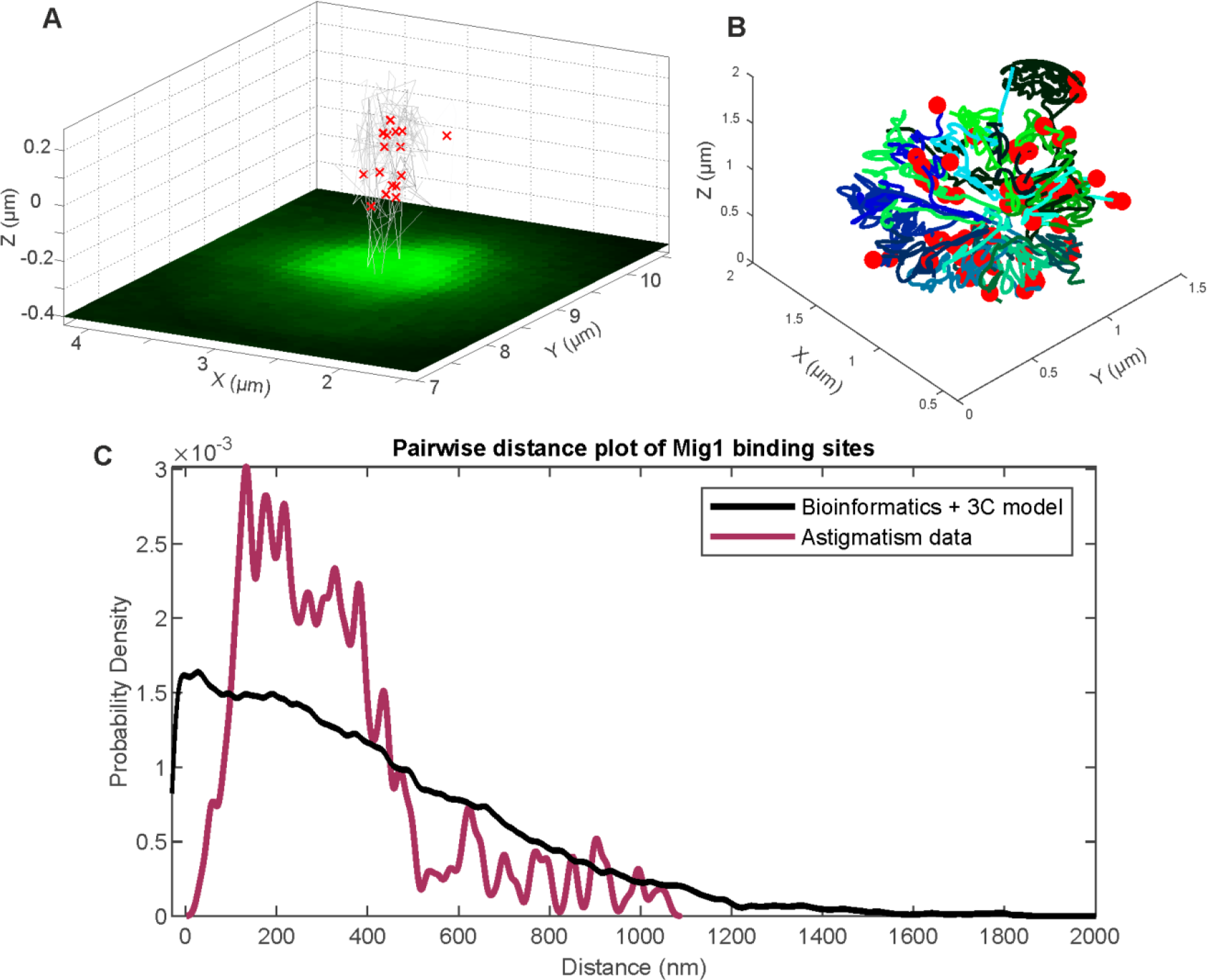
A. Mig1-GFP in a single live yeast cell (green) with the 3D position of immobilized Mig1 molecules in the nucleus marked as red crosses and their trajectories indicated as grey lines. B. The 3C model of yeast chromosomal DNA (green and blue lines) from reference [36]) with the position of Mig1 binding sites within promoters marked as red dots from reference [7]). C. The distribution of pairwise distances of detected immobilized Mig1 foci in the nucleus (maroon) and predicted from Mig1 binding sites in promoters in the 3C model (black).

We took the 3D centroid positions of immobile Mig1 tracks as the putative positions of Mig1 binding sites in the genome and an indicator of 3D genome architecture (Fig. 4A). We compared these positions to those obtained from a predictive model, using bioinformatics to map out likely Mig1 binding sites within promoters onto a 3C model of yeast chromosomal DNA (Fig. 4B) [7]. Potential Mig1 binding sites were identified by analyzing the whole genome of *S. cerevisiae* in order to find DNA sequences that fit a consensus Mig1 target generated based on 14 known Mig1 binding site sequences [37]. To compare experimental and modeling outcomes, we calculated the distribution of pairwise distances of observed immobile Mig1 foci in 25 cells and of potential Mig1 binding sites in the model (Fig. 4C). The distribution from the astigmatism data is different to the theoretical prediction. The mean pairwise separation of the predicted distribution is 417 ± 30 nm, ±SE, compared against 330 ± 7 nm, with a Student’s *t*-test indicating different means (*P*<0.0001), although if pairwise distances greater than our working 1 μm range are excluded from the predicted distribution, the mean is 368 ± 25 nm which although marginally closer to our experimental measurements is still statistically different (*P*<0.0001),.

This intriguing observation may result from several possibilities. For example, if only certain areas of the genome are undergoing active transcription, then it is possible that the range of the pairwise differences could be relatively higher than expected but that the peak value could potentially be smaller if the regions of active transcription are themselves relatively clustered. Similarly, the clustered nature of Mig1 implies multivalency of binding to DNA, and this in turn may result in condensation effect of separate DNA strands which are linked by a cluser, thus shifting the mean pairwise distance. It may also be the case that only a subset of spatially clustered Mig1-regulated genes have Mig1 bound to the target promoters at any one time: a subset of Mig1 clusters might also bind transiently (at least over a duration of four consecutive image frames or 20 ms used for the diffusion coefficient estimates), but in a relatively immobile state, to regions of the genome which are not specific promoter regions. Such a phenomenon could indeed be functionally important in intersegmental transfer of clusters between DNA segments [38]. This putative hopping motion may reduce the search time for Mig1 to ultimately find its gene targets, and thus might be expected to occur at regions which are not collocated with the Mig1 binding sites themselves. Hopping between separate DNA segments may of course result in transient immobility to the translocation along the original DNA segment prior to reaching the destination DNA segment.

Either way, our 3D imaging approach has very clearly revealed significant tiers of complexity to the dynamic architecture of the genome, which slower, less precise and less physiologically relevant techniques would simply fail to render. This insight alone is incredibly valuable. Of course, it remains to be determined in future studies precisely what are the key explanations for this heterogeneity in the genomic architecture, but our methodology shows promise in being able to enable such future insights. While the 3C model is an ensemble technique which represents an average conformation of a genome, our method reconstructs 3D genome architecture at a given time and thus enables truly detailed studies of its dynamics in a single living cell.

## Conclusions

Here we describe a method which enables us to map out the 3D positions of immobile fluorescent DNA-binding molecules, which thereby act as a valuable proxy to indicate the genome architecture in live yeast cells. We apply it to the Mig1 transcription factor for which we have a predicted model of the 3D position of binding sites within the genome from a knowledge of its binding sequences within the target gene promoters applied to prior 3C data. This allows us to access information about the functional 3D genome architecture in live cells. Recent methods now allow single-cell 3C to be combined with microscopy [39], although currently only using lateral imaging (i.e. 2D, in 2D spatial dimensions). If this technique were combined with our 3D method, it could potentially unlock transformative levels of information about genome architecture. Our method here is timely, given the increasingly revealed complexity in the dynamics of transcription factors and the impact of 3D DNA geometry on gene regulation [40], as well as other new methods using single-molecule FISH (smFISH) approaches which can map chromatin in 3D at a single-cell level [41], [42]. Our general method could in principle be extended to many other native proteins or artificially expressed markers for specific genome loci in both eukaryotic and prokaryotic organisms, such as tagging of the *Lac* operon of *Escherichia coli* [43], [44] which could then be used to report on the 3D architecture of highly specific sites within the prokaryotic genome. Although as it stands the spatial precision of our 3D imaging toolkit is poor compared to 3C variant approaches and smFISH there is a substantive advantage in our approach in enabling us to interrogate the genomic 3D architectures using very high time resolution of milliseconds on single live cells. A really exciting future of this technology could perhaps lie with combining this exceptional time resolution with a higher spatial resolution technology such as 3C or smFISH based approaches.

## Supplementary Information

**Supplementary Figure 1:**
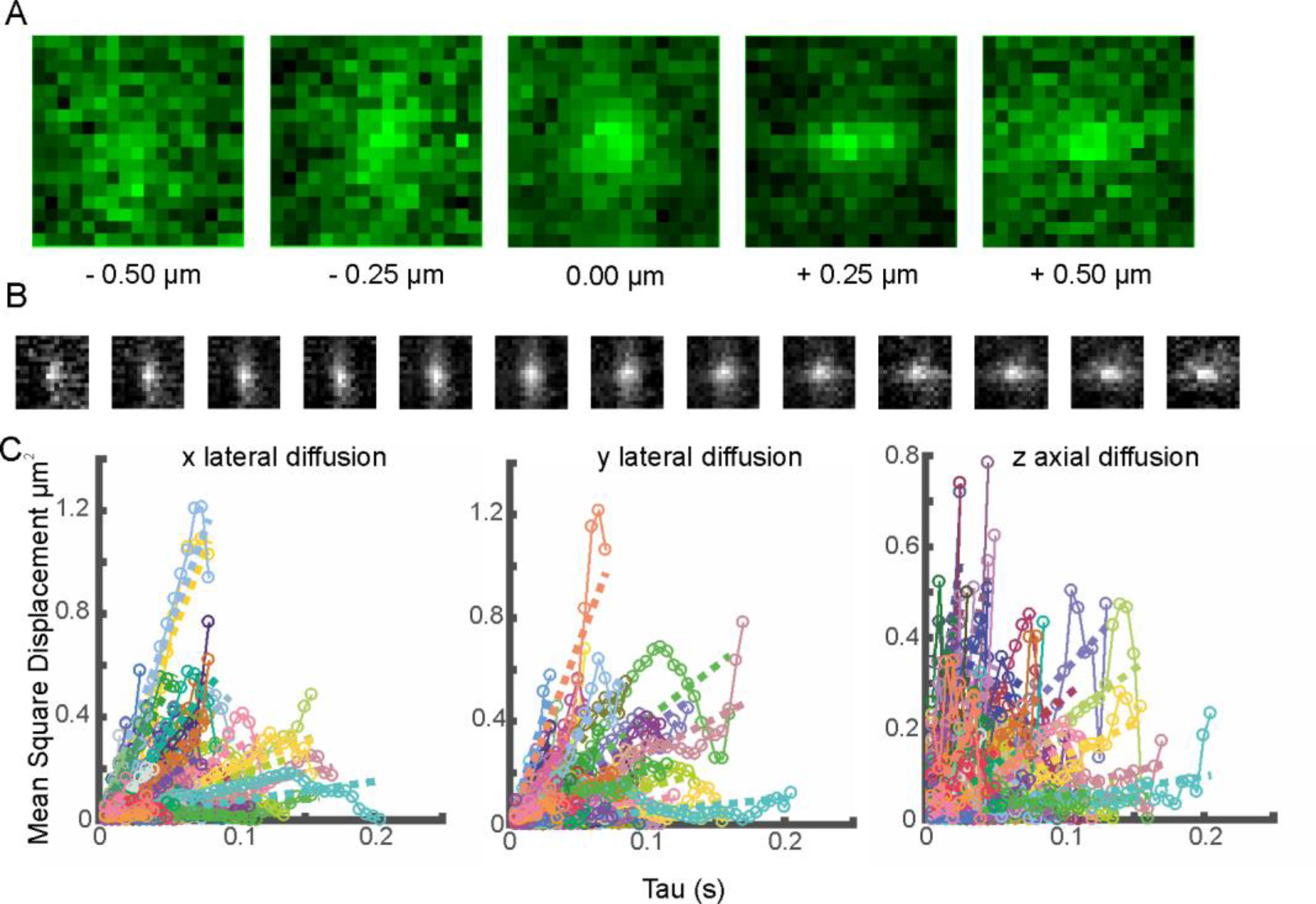
*A. Mean PSF as a function of axial distance z as based on in vitro GFP images. B. Interpolated PSF as a function of z used in simulations. C. Mean square displacement vs. time interval (Tau) for simulated trajectories*

